# Involvement of Mitophagy in Endothelin-1 Mediated Neurodegeneration in Rodent Models of Glaucoma

**DOI:** 10.64898/2026.07.02.735939

**Authors:** Calvin D Brooks, Bindu Kodati, Sanjna Prasad, Jacob Cunningham, Poojan Patel, Michael Mangan, Stacy Curry, Drayanna K FoxRun, Aysha Ehsan, Ojasvi Arya, Hayden Flume, Kishor Kunwar, August E Woerner, Denise M Inman, Dorota L Stankowska, Raghu R Krishnamoorthy

## Abstract

The ultimate cause of blindness in glaucoma is the death of retinal ganglion cells, and understanding the mechanism behind retinal ganglion cell loss during glaucoma could lead to the development of novel treatments for glaucoma. Endothelin-1 has been shown to mediate retinal ganglion cell death during glaucoma through impairment of mitochondrial function. Retinal ganglion cells are highly metabolically active, and susceptible to oxidative damage and decreased respiratory capacity. Mitophagy is the process whereby damaged mitochondria are degraded to prevent further propagation of oxidative damage. The current study evaluates the effect of endothelin-1 on mitophagy in retinal ganglion cells. Electron microscopy revealed endothelin-1 administration lead to a decrease in healthy mitochondria in the optic nerve. The MitoQC mouse was used to evalute mitophagy in response to endothelin-1, along with immunohistochemical analysis of mitophagy proteins. Mitophagy follows different trends in the optic nerve and retinal ganglion cell bodies following endothelin-1 administration, mitophagy was increased in the optic nerve but decreased in the retina following endothelin administration. With elevation of intraocular pressure, mitophagy was increased in the retina but decreased in the optic nerve. In retinal ganglion cells, parkin expression and activation was unchanged 24 hours after endothelin-1 administration, but was decreased 72 hours following endothelin-1 administration. Taken together, these results suggest that endothelin-1 impacts mitophagy through parkin-independent mechanisms in retinal ganglion cell bodies, and the ganglion cell bodies and optic nerve appear to have different responses to endothelin-1.

## 1 Introduction

Glaucoma is a disease characterized by degeneration of the optic nerve, thinning of the inner retina, and death of retinal ganglion cells, leading to vision loss that typically starts in the periphery and progresses centrally (1,2). While there are many risk factors associated with the increased prevalence or incidence of glaucoma, the main risk factor that can predict the progression of glaucoma is elevated intraocular pressure (IOP) (3). Current treatment strategies focus on lowering intraocular pressure, resulting in varying degrees of success (4). Several classes of drugs have been approved for lowering of intraocular pressure, with the most recent being nitric oxide donors and rho kinase inhibitors (5). In addition to drug treatment, surgical interventions are very common, including trabeculoplasty, trabeculectomy, shunts and devices to increase aqueous humor drainage, as well as cyclophotocoagulation of the ciliary body to decrease aqueous humor formation (4). Even in normotensive glaucoma, characterized by glaucomatous optic neuropathy in the absence of an elevation in intraocular pressure, the current treatment strategy is to lower IOP below the physiologic values (6). However, these IOP-lowering treatments can also lose effectiveness over time (7–9). Long term success of IOP lowering treatments in pediatric glaucoma leaves a lot to be desired (10–14). While the effectiveness of IOP-lowering treatments is of concern, even when IOP is reduced to below physiologi levels in normotensive glaucoma, glaucomatous damage still progresses (6). These factors combined lead to worse patient outcomes. In one study, over 30% of patients became blind in at least one eye within 20 years of diagnosis with glaucoma (15). Better clinical outcomes may be achieved by developing neuroprotective strategies to prevent retinal ganglion cell degeneration regardless of IOP (6,16,17).

Endothelin-1(ET-1) is a small vasoactive peptide, known for its vasoconstrictive activity and role in the pathogenesis of several cardiovascular diseases (18–23). The 3 isoforms of endothelin (ET-1, ET-2, ET-3) and both endothelin receptors (ET_A_, and ET_B_) are expressed throughout the eye including in the retina (24,25). The major sources of endothelins within the eye are the non-pigmented ciliary body epithelium, retinal pigment epithelium and optic nerve head astrocytes (26–32). ET-1 was found to be increased in the plasma and aqueous humor of primary open angle glaucoma patients, and in the aqueous humor of dogs with glaucoma (33,34). ET-1 was increased in the aqueous humor and optic nerve head in rats subjected to the Morrison’s model of experimentally induced ocular hypertension (35,36). ET-1 levels were also increased in an IOP-independent manner, and ET-1 increases are also seen the aqueous humor of patients with normotension glaucoma (37,38). During glaucoma, astrocytes become a prominent source of ET-1 in the retina and optic nerve (30–32). In addition to secreting ET-1, astrocytes also respond to ET-1 by increasing activation, migration, and proliferation, and ET-1 can also induce pathologic remodeling of the lamina cribrosa and optic nerve head region (25,30–32,35,39–43). Increases in ET-1, independent of IOP increases, contributes to optic neuropathy and retinal ganglion cell (RGC) death (25,31,32,42–46). In addition to astrogliosis, ET-1 can also induce hypoxic injury in RGCs (25,46,47). ET-1 also inhibits axonal transport in RGCs (44). Endothelin receptor antagonists protected RGCs in a rat model of glaucoma, despite having no effect on IOP, demonstrating this class of drug as a potential neuroprotective treatment for glaucoma (48).

Endothelin receptors are also expressed on retinal ganglion cells, and their expression increases during ocular hypertension and following administration of ET-1 (45,46,49,50). Activation of endothelin receptors on primary cultured RGCs leads to cell death (45). The absence of vascular effects in cultured RGCs reveals that ET-1 directly mediates cellular injury, the mechanisms of this pathway are still being explored. Many changes in RGC gene expression are seen in response to ET-1, increases in several genes in the Bcl-2 family may be the cause of cell death in response to ET-1 (50). The expression of several mitochondrial genes are also changed in RGCs following ET-1 treatment (51). Most notably is a reduction in ATP synthase, and it corresponds to decreases in oxygen consumption rate (51). The decrease in ATP synthase is also seen in Morrison’s model of ocular hypertension (51). Endothelin treatment produced sustained reductions in baseline and maximal oxygen consumption rate, and increases in reactive oxygen species, in cultured RGCs (52). ET-1 causes similar mitochondrial dysfunction in other neuron populations and has been implicated in hippocampal neuron death during Alzheimer’s disease (53).

Schrier and Falk (2011) describe the mitochondria as the powerhouse of cells and mitochondrial dysfunction is implicated in the etiology of several eye disorders, including, diabetic retinopathy, age-related macular degeneration and glaucoma (54). Mitochondria are particularly important for any metabolically active cells, including output neurons in the central nervous system, hence, mitochondrial dysfunction is implicated in several neurodegenerative diseases (55). Mutations in mitochondrial DNA are responsible for eye diseases like Leber Hereditary Optic Neuropathy (54). Even with no direct genetic cause, mitochondrial dysfunction plays a causal role in complex retinopathies like age-related macular degeneration (56–59). Retinal ganglion cells, having long axon projections with the proximal end unmyelinated until the post-laminar region of the optic nerve head, have high metabolic demand, and are therefore quite susceptible to mitochondrial dysfunction and oxidative stress (60). Decreases in mitochondrial function and metabolism contribute to cell death and axonal degeneration in glaucoma (61–64). Oxidative stress has been observed in RGCs during glaucoma, increasing reactive oxygen species and inducing oxidative damage (65–67). Furthermore, antioxidant treatments protect RGCs during experimentally induced ocular hypertension (65,68,69). In addition, mitochondrial dysfunction in glaucoma can promote inflammation, which further mediates cell death (70).

Mitophagy is a specialized form of autophagy whereby damaged mitochondria are degraded and their contents recycled (71–74). Working in concert with mitochondrial fission, fusion, and biogenesis, mitophagy is an important process for ensuring adequate bioenergetics while maintaining quality control of mitochondria, removing damaged mitochondria that produce excess reactive oxygen species (73,75,76). Impairments in mitophagy have been implicated in many neurodegenerative diseases (77–79). A recent review by Brooks et al. (2023) covers findings on mitophagy in common retinal neurodegenerative diseases (79). Briefly, mitophagy is known to have a pathogenic role in age-related macular degeneration and diabetic retinopathy (80–82). The role of mitophagy in glaucoma has not been fully elucidated. Some researchers report increased mitophagy during glaucoma, while some report mitophagy decreases(83–86). Discrepancies may be due to differences between models or in measurements used. Obanina et al (2022) found significant mitochondrial damage in glaucoma patients, but also found evidence of increased mitophagy, these results are similar to those seen by Zeng et al. (2020) in glucocorticoid-induced ocular hypertension in mice (83,84). However, others have found that increasing mitophagy is neuroprotective in glaucoma (85). It is possible that mitophagy does increase, but not to the level needed to provide adequate protection of cells. Gurubaran et al. (2020) found that their mouse model of age-related macular degeneration had increased expression of PINK1 and Parkin, common initiators of mitophagy, but there was a lack of an increase in autolysosome formation, presumably creating a bottleneck in the mitophagy pathway (86). Alternatively, differences in outcomes could represent temporal changes in mitophagy following IOP elevation, Ma et al. (2025) found that mRNA expression of parkin, optineurin, LC3, and LAMP1 were increased 3 days following LASER photocoagulation of the trabecular meshwork to induce IOP elevation, but expression of all but LC3 tapered off, and was reduced at 14 days compared with control rats (87). Further, their neuroprotective treatment increased the expression of all the aforementioned markers at 14 days, suggesting that increasing mitophagy promotes the long-term survival of RGCs (87). Maddineni et al. (2026) also found decreased mitophagy in RGCs following 5 weeks of dexamethasone-induced ocular hypertension in mice (88). Evaluating the expression of proteins involved in mitophagy is a common method of evaluating mitophagy, but the results can be hard to interpret at times (79). Our lab found that ET-1 decreased the colocalization of lysotracker and mitotracker in cultured RGCs, and decreased colocalization of Tom20 and LC3B in vivo (52). A similar trend of decreased colocalization of Tom20 and LC3B was seen in Morrison’s model of ocular hypertension in rats (52). This provides preliminary evidence that ET-1 may inhibit mitophagosome formation in RGCs during glaucoma, and could be a novel mechanism of RGC loss in glaucoma (52). The purpose of the current study was to further evaluate mitophagy and mitochondrial health and identify changes in key components of the mitophagy machinery in RGCs exposed to ET-1.

## 2 Methods

### 2.1 Animals

All animal experiments were carried out in accordance with the policies of the Association for Research in Vision and Ophthalmology (ARVO) for use of animals in research and approved by the Institutional Animal Care and Use Committee (IACUC) (animal protocol #IACUC-2023-0024) at the University of North Texas Health Science Center at Fort Worth, TX, USA. Animals were housed in standard conditions unless otherwise stated. MitoQC mice (C57Bl/6-Gt(ROSA)26Sor^tm1(CAG-mCherry/GFP)Ganl^) (age 3 months or 1 year) were bred in house. We thank Ian Ganley for providing the mutant mouse line (Allele: Gt(ROSA)26Sor<tm1(CAG-mCherry/GFP)Ganl>), INFRAFRONTIER/EMMA (www.infrafrontier.eu, PMID: 25414328), and EMMA node at Mary Lyon Centre at MRC Harwell from which the mouse line was distributed (RRID:IMSR_EM:11343). The MitoQC mouse uses a dual-flourophore labelling system for the detection of mitophagy in real time (89,90). For one set of experiments, C57BL6/J mice (the parent strain of the MitoQC mouse) (age 3 months) were obtained from Jackson Labs (Bar Harbor, ME).

#### 2.1.1 Injection of Endothelin-1

MitoQC Mice (adult and aged) received bilateral intravitreal injection of ET-1 (1 nmole) or vehicle (water). Injections were performed under isoflurane anesthesia. At three timepoints, 24, 48, or 72 hours following injection, mice were anesthetized using an intraperitoneal (IP) injection of pentobarbital, followed by euthanasia via intracardiac injection of pentobarbital, and confirmed by cervical dislocation. One eye from each animal was collected for live (unfixed) imaging, as described below. The other eye was enucleated and fixed in 4% paraformaldehyde (PFA) (Electron Microscopy Sciences, Hatfield PA) for paraffin embedding and sectioning. One optic nerve from each mouse was fixed in 4% PFA for cryosectioning.

In a separate experiment, C57Bl6/J mice received an intravitreal injection of ET-1 or vehicle in the same manner and at three time points, 24 hours, 72 hours, or 7 days following injection, mice were euthanized using the same procedure. Both eyes were fixed in 4% PFA, one eye was dissected and the retina stained and mounted flat to quantify retinal ganglion cell loss, the other eye was embedded in paraffin for sagittal sectioning. Additionally, one optic nerve from each mouse was fixed in ½ strength Karnovsky’s fixative and processed for electron microscopy as described below.

#### 2.1.2 Microbead Model of Ocular Hypertension

In an effort to determine the effect of elevations in intraocular pressure on mitophagy, MitoQC mice received a unilateral intracameral injection of magnetic microbeads to raise intraocular pressure as described previously (91). Injections were performed under isoflurane anesthesia, and animals were housed in constant dim light (90 lux) before and after the procedure. Intraocular pressure was monitored under anesthesia twice a week before and after microbead injection, and following 1 week of IOP elevation, IOP exposure was calculated as the area under the IOP elevation curve, total IOP exposure >16 mmHg*Days was the criterion for inclusion in the study, representing mild IOP increases consistent with the early stages of glaucoma. Both eyes and optic nerves from these mice were fixed in 4% paraformaldehyde, eyes were paraffin embedded for sagittal sectioning while the optic nerves were cryosectioned.

### 2.2 Quantitation of retinal ganglion cell death

One eye from C57Bl6/J mice were used to quantify RGC death following ET-1 injection. Eyes were fixed in paraformaldehyde and stored in PBS, the eye was dissected and retina transferred to a 48 well plate for immunostaining. Retinas were treated with PBS containing 0.3% Triton-X100 for 1 hour with the solution changed every 20 minutes, then blocked with 10% normal donkey serum with 0.3% Triton-X100 for 1 hour. Retinas were stained with a rabbit anti-RBPMS antibody (GeneTex, Irvine, CA; #GTX1186, 1:250) in 10% normal donkey serum with 0.3% Triton X-100 for 3 days at 4°C. Retinas were washed for 3 hours in PBS, with PBS changed every 30 minutes, and incubated overnight at 4°C in a solution of Alexa-flour conjugated donkey anti rabbit (Invitrogen, Waltham MA) secondary antibody in PBS containing 0.1% Triton-X100. Retinas were washed to remove excess antibody, spread flat on a microscope slide and sealed with a coverslip. Retinal flat mounts were imaged on a Keyence BZ-X710 fluorescent microscope. Two images were taken in each of 3 eccentricities within each of 4 quadrants of the retina. The cell count for each image was obtained using an automated cell counter in FIJI (ImageJ), and for each animal the count was averaged separately for each eccentricity.

### 2.3 Transmission electron microscopy of the optic nerve

C57BL6/J mice were injected with either ET-1 or vehicle for 24 hours, 72 hours, or 7 days, following which one optic nerve was collected for transmission electron microscopy to analyze mitochondrial morphology. After dissection, optic nerves from the mice were fixed in ½ strength Karnovsky’s fixative for more than 3 days.

Nerves were washed in 0.1M sodium cacodylate buffer, stained with a solution of 1% osmium tetroxide with 1.5% potassium ferrocyanide in cacodylate buffer, and washed again with cacodylate buffer. Nerves were then treated with 2.5% uranyl acetate, then dehydrated in a graded series of ethanol solutions, then propylene oxide, then finally the nerves were embedded in eponate (Ted Pella, Redding, CA), which was allowed to cure overnight at 70°C.

Once embedded, optic nerves were sent to the Central Microscopy Research Facility at the University of Iowa for ultramicrotome sectioning and transmission electron microscopy on a Hitachi H-7800, AMT NanoSprint15 high-resolution, high-sensitivity camera system with 5056 x 2960 pixel sensor. Optic nerve sections were split into 5 zones (central, left, right, superior, inferior) and two images were taken in each zone at 10,000X magnification. Areas were selected for imaging based on a high density of axons, this was to normalize the number of axons in each image and ensure enough axons could be sampled to determine the mitochondrial density of these axons.

Imaging and analysis were performed in a masked manner. Mitochondria within optic nerve axons were counted and scored 1-5 based on the appearance of the cristae (with 1 being the worst and 5 being the best). This is an accepted proxy for mitochondrial health as mitochondria without well-defined, dense, organized, and regularly spaced cristae have limited ability to perform oxidative phosphorylation (92–95). Mitochondrial health scores and the number of mitochondria per image were averaged for each animal and compared across treatment groups and time points.

### 2.4 Immunostaining of retinal sections

Eyes for sagittal sectioning were fixed in 4% paraformaldehyde for 4 hours at room temperature, then washed in 70% ethanol and stored at 4°C in 70% ethanol before paraffin embedding. An automated tissue processor (Tissue-Tek, Nagano, Japan) was used to dehydrate the eyes in a graded series of ethanol concentrations, followed by xylene (Mercedes Scientific, Lakewood Ranch, FL) and paraffin (Mercedes Scientific, Lakewood Ranch, FL). Eyes were then embedded in paraffin blocks using a Leica tissue embedding station (Leica, Wetzlar, Germany), and sectioned using a rotary microtome (Leica, Wetzlar, Germany).

Slides containing sagittal sections of the mouse eye were deparaffinized in xylene and rehydrated using a graduated series of ethanol solutions and ending with phosphate-buffered saline (PBS). Sections were permeabilized with a solution of 0.1% sodium citrate and 0.1% Triton X-100, then nonspecific binding sites blocked with buffer containing 5% bovine serum albumin (BSA) (Sigma Aldrich, St. Luis, MO), and 5% normal donkey serum (Jackson Immunoresearch, West Grove, PA) in PBS for 1 hour. Slides were incubated in a solution of specific primary antibodies overnight at 4°C, excess antibody was washed off before staining with corresponding secondary antibodies for 90 minutes at room temperature. Slides were then counter-stained with 4’,6-diamidino-2-phenylindole (DAPI) (Sigma Aldrich, St. Luis, MO) to identify nuclei and mounted using DEPEX (Electron Microscopy Sciences, Hatfield PA).

MitoQC eyes were only immunolabeled with an anti RBPMS antibody (Thermo Fisher, Waltham, MA; #PA5119676, 1:200) and an appropriate secondary antibody that did not overlap with endogenous fluorescent signal (Jackson Immunoresearch Donkey anti Guinea Pig Alexa Flour 647; 1:1000). Sections were imaged in a Keyence BZ-X710 fluorescent microscope. The ganglion cell layer was identified by the presence of RBPMS positive cells, mitophagy was quantified within this layer using the procedure established by Montava-Garriga (2020) (90).

C57Bl6/J eyes were immunolabeled with an anti RBPMS antibody (Thermo Fisher, Waltham, MA; #PA5119676, 1:200) as well as antibodies targeting proteins involved in mitophagy. Antibodies used include Tom20 (Sigma Aldrich, St. Luis, MO; # WH0009804M1-100 1:50), Lamp1 (Novus Biologicals, Centennial, CO; #NB120-19294, 1:200), LC3B (Sigma Aldrich, St. Luis, MO; #L7543, 1:200), Parkin (Thermo Fisher, Waltham, MA; #390900, 1:100), phospho-Ser65-Parkin (Thermo Fisher, Waltham, MA; #PA5114616, 1:200). Slides were then tagged with alexa-fluor conjugated secondary antibodies (Thermo Fisher, Waltham, MA; or Jackson Immunoresearch, West Grove, PA), and sections were imaged in a Keyence BZ-X710 fluorescent microscope. Fluorescence intensity (as measured by integrated density) was calculated in a region of interest covering the ganglion cell layer and nerve fiber layer of the retina. Blank corrected fluorescent values were compared between groups. Additionally, Manders’ correlation coefficients M1 and M2 were used to assess colocalization of certain markers.

### 2.5 Mitophagy in MitoQC Optic Nerves

Optic nerves from MitoQC mice, following intravitreal injection of ET-1 or vehicle, or following IOP elevation, were fixed in 4% PFA for 4 hours, washed with phosphate buffered saline (PBS) (Teknova, Hollister, CA) and stored at 4°C until embedding. Optic nerves were washed in increasing concentrations of sucrose, before embedding in cryopreservation media (Fisher Scientific, Waltham, MA) and freezing at -80°C. Sagittal sections of frozen optic nerves were obtained using a cryostat (Leica, Wetzlar, Germany) and affixed to slides. Sections were washed with PBS to remove cryopreservation media, and a coverslip affixed using DEPEX mounting media (Electron Microscopy Sciences, Hatfield, PA) before imaging in a Keyence BZ-X710 Fluorescent microscope, 40X images were taken at different points along the proximal optic nerve.

Images were analyzed using a procedure adapted from Montava-Garriga et al (2020) (90). mQC_counter.ijm was edited to enable auto thresholding using “Huang dark” thresholding method on the red channnel, the selection was created from the threshold image and added to ROI manager to fulfil the requirement for processing “ROIs for cells *only* should be in ROI manager” in original mQC_counter macro. This modification allowed quantification of mitophagy across the whole image.

### 2.6 MitoQC retina live imaging

Following intravitreal injection of ET-1 in MitoQC mice, one eye was enucleated and unfixed retinal flat mounts were prepared for live imaging of mitophagy in RGCs. The unfixed retina was dissected, and mounted flat with the RGCs superior on a microscope slide with PBS containing 10% glycerol. The slide was immediately imaged on a Zeiss LSM 880 (Zeiss) with Airyscan detector and processed with Airyscan processing in Zeiss Zen 2.3. Images were acquired using a 63X /1.4NA oil immersion objective with pixel size of 39nm. Excitation laser of 488nm and 561nm with emission filter BP495-550_+ LP570 was used to acquire image. C<u>ells</u> were identified by characteristic shape, size, and morphology, and each image contained roughly a single cell. Additional wide FOV images were taken but not used in analysis. The RGC layer was determined by proximity to the inner retina, and Z-stacking was used to cover the entire ganglion cell layer and nerve fiber layer. Images were analyzed using the same procedure as above (see section 2.5).

### 2.7 Statistical analysis

All microscopy images were analyzed in FIJI (ImageJ) in a masked manner (96). Standard tools and plugins were used with the exception of the mitoQC counter macro as described above (90). Data gathered from each image was averaged per animal/eye and imported into Prism (Graphpad Software, Boston, MA) for statistical analysis. If outliers were suspected, the ROUT method with Q=1% was used to identify outliers. Normality and homoscedasity was assessed for every dataset. For analysis of groups across multiple factors (e.g. treatments tested at multiple time points), a full-effects model two-way ANOVA with Fisher’s LSD post-hoc, or multiple Mann-Whitney tests with Holm-Sidak correction as a non-parametric alternative. If only two groups are being compared, a T test or Wilcoxon Rank-Sum test was used. For IOP-elevated mice, the IOP elevated eye/optic nerve was compared with the contralateral eye/optic nerve using a Wilcoxon paired signed rank test. All graphs represent mean ± SEM with the individual data points overlayed for each eye/animal. MitoQC Live imaging data was imported into R for further visualization and exploratory analysis utilizing the GGally and Tidyverse R packages (97,98).

## 3 Results

### 3.1 Intravitreal ET-1 Administration Causes RGC Loss in C57BL6 mice

Intravitreal injection of ET-1 in C57BL6/J mice led to significant RGC death in the central retina 72 hours following injection (p=0.01), and slightly more cell death was detected 7 days following injection (p=0.0054) (Figure 1). The same trend was significant in the mid-peripheral retina (p=0.02 and p=0.04 respectively), but was not significant in the peripheral retina (Supplemental Figure 1).

**Figure 1:**
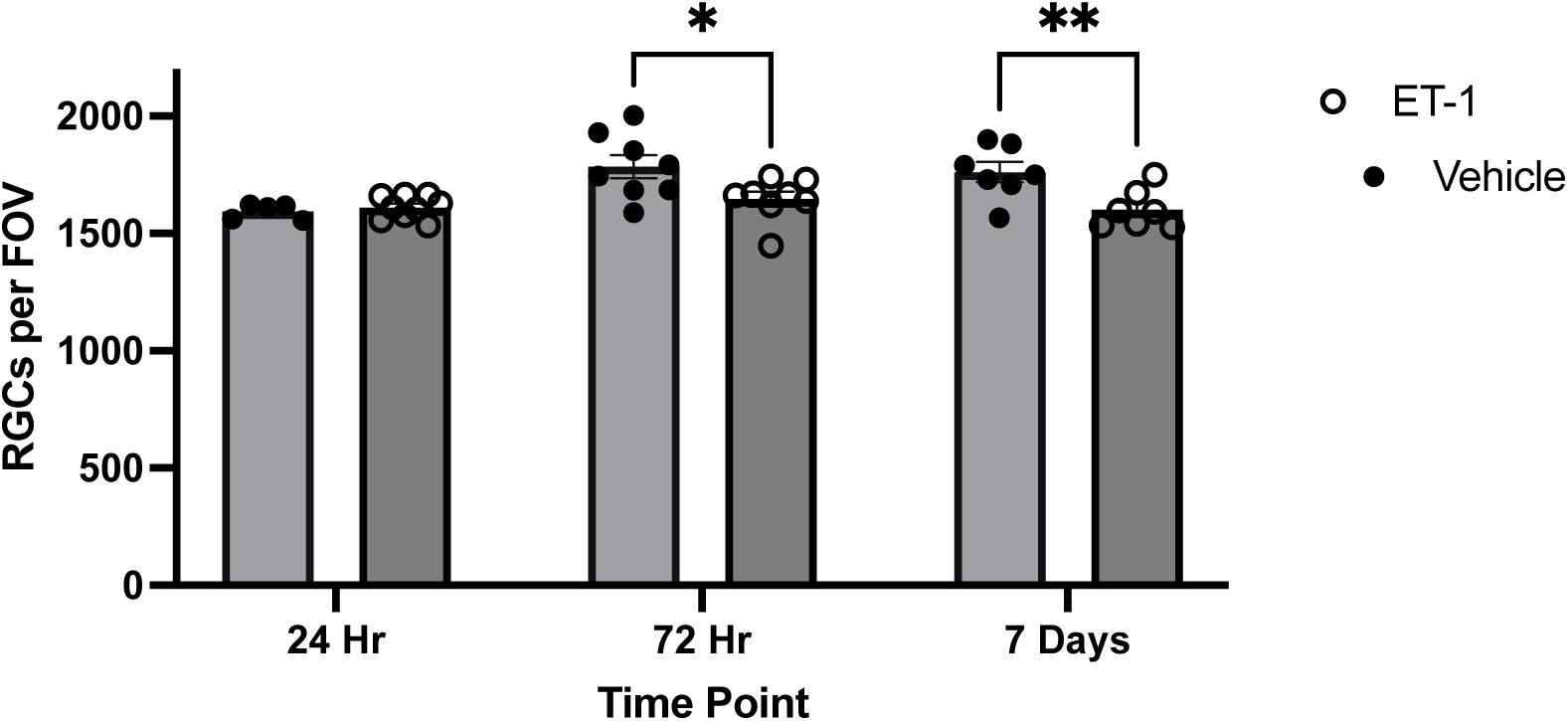
RGC Loss Occurs 72 hours following ET-1 Injection: C57BL6/J mice received IVT injection of ET-1 (1 nMole) or vehicle, cell death was evaluated through the quantification of RBPMS positive cells per field of view in retinal flat mounts. Significant cell death was seen 72 hours, and 7 days following ET-1 injection in the central retina (p=0.01 and p=0.005 respectively, two-way ANOVA with Fisher’s LSD).

### 3.2 ET-1 Disrupts Mitochondrial Dynamics in the Optic Nerve

To further examine the effect of ET-1 administration on mitochondrial health, cross sections of optic nerves from C57BL6/J mice injected with ET-1 or vehicle were imaged using transmission electron microscopy. ET-1 decreased the number of mitochondria in the optic nerve (p=0.02, two-way ANOVA treatment effect). In post-hoc testing, the difference is only significant 24 hours following injection (p=0.05), suggesting this decrease in mitochondria recovers over time following injection (Figure 2a). A two-way ANOVA revealed a significant effect of time on mitochondrial health following IVT injection (p=0.04). Mitochondrial health was reduced 24 hours following IVT injection in both groups, and increased over time. However, in post-hoc tests the effect was only significant in mice injected with ET-1 (p=0.04) (Figure 2b). This suggests a difference between ET-1 and vehicle injected mice may exist 24 hours following injection that we are not sufficiently powered to detect.

**Figure 2:**
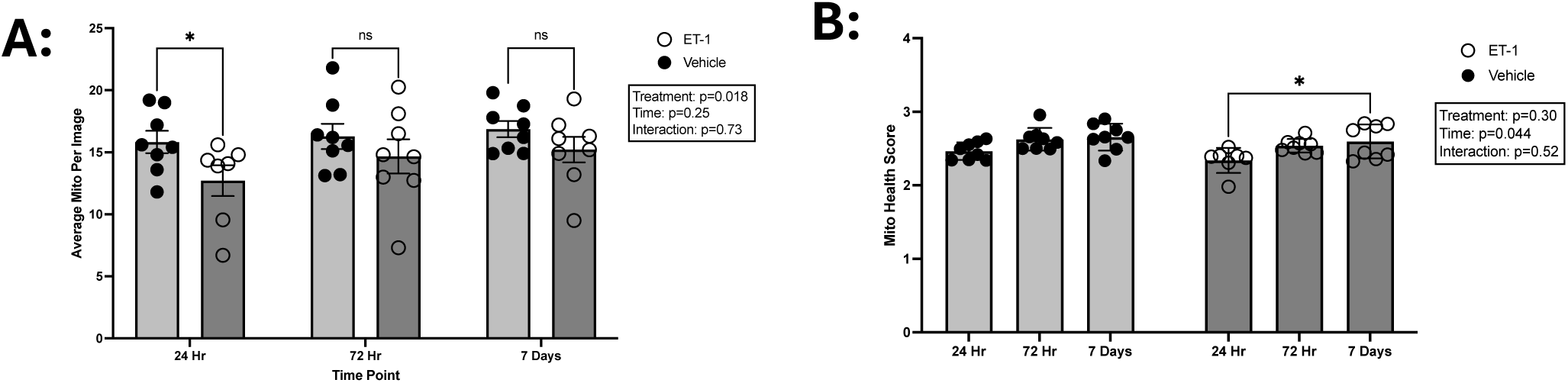
Mitochondrial Morphology In Axons Affected By ET-1. **A:** Transmission electron microscopy revealed mitochondrial density in RGC axons is reduced following ET-1 injection (p=0.05, two-way ANOVA with Fisher’s LSD). **B:** Mitochondrial cristae structure is disrupted following IVT injection of ET-1 and vehicle, but significant recovery of cristae structure is seen following ET-1 injection (p=0.04, two-way ANOVA with Fisher’s LSD).

### 3.3 ET-1 Affects Expression of Mitophagy Proteins

Immunostaining of TOM20 and LC3B in sagittal sections of C57BL6/J mouse eyes injected with ET-1 or vehicle was used to assess autophagosome formation. Mann Whiteny tests with Holm-Sidak correction revealed that the expression of both TOM20 and LC3B were reduced 24 hours following ET-1 injection (adjusted p=0.002 and p=0.004 respectively) (Figure 3a). In separate sections of the same animals, TOM20 and LAMP1 were stained to evaluate mitolysosome formation. A two-way ANOVA revealed a significant time effect on LAMP1 expression (p=0.05), but no individual relationships were significant in post-hoc testing (Figure 3a). No significant differences were detected in Mander’s correlation coefficient (M1 or M2) between TOM20 and LC3B or between TOM20 and LAMP1 (two-way ANOVA), although colocalization of TOM20 and LAMP1 was generally lower than colocalization of TOM20 and LC3B, which was expected (Figure 3b).

**Figure 3:**
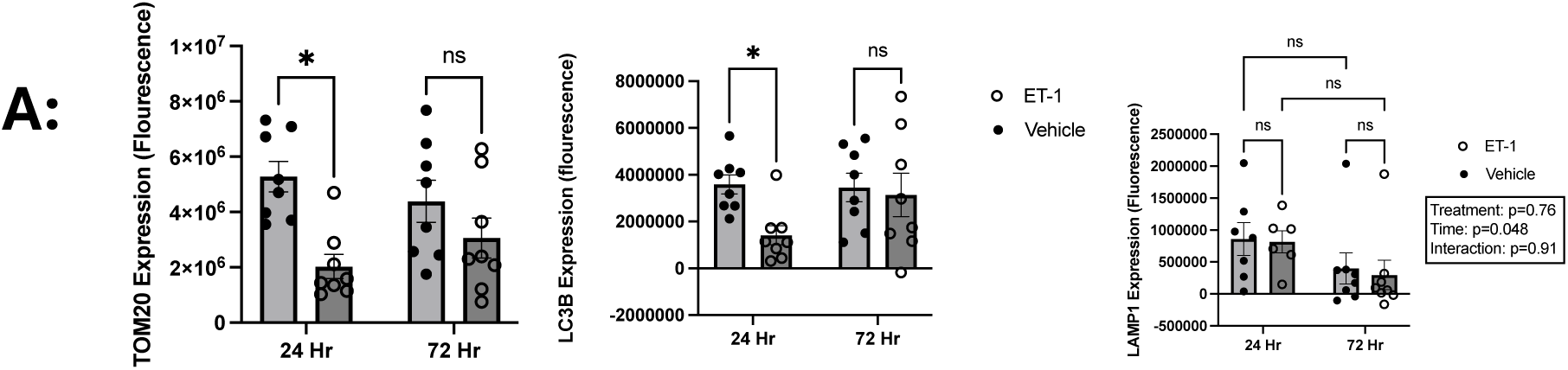
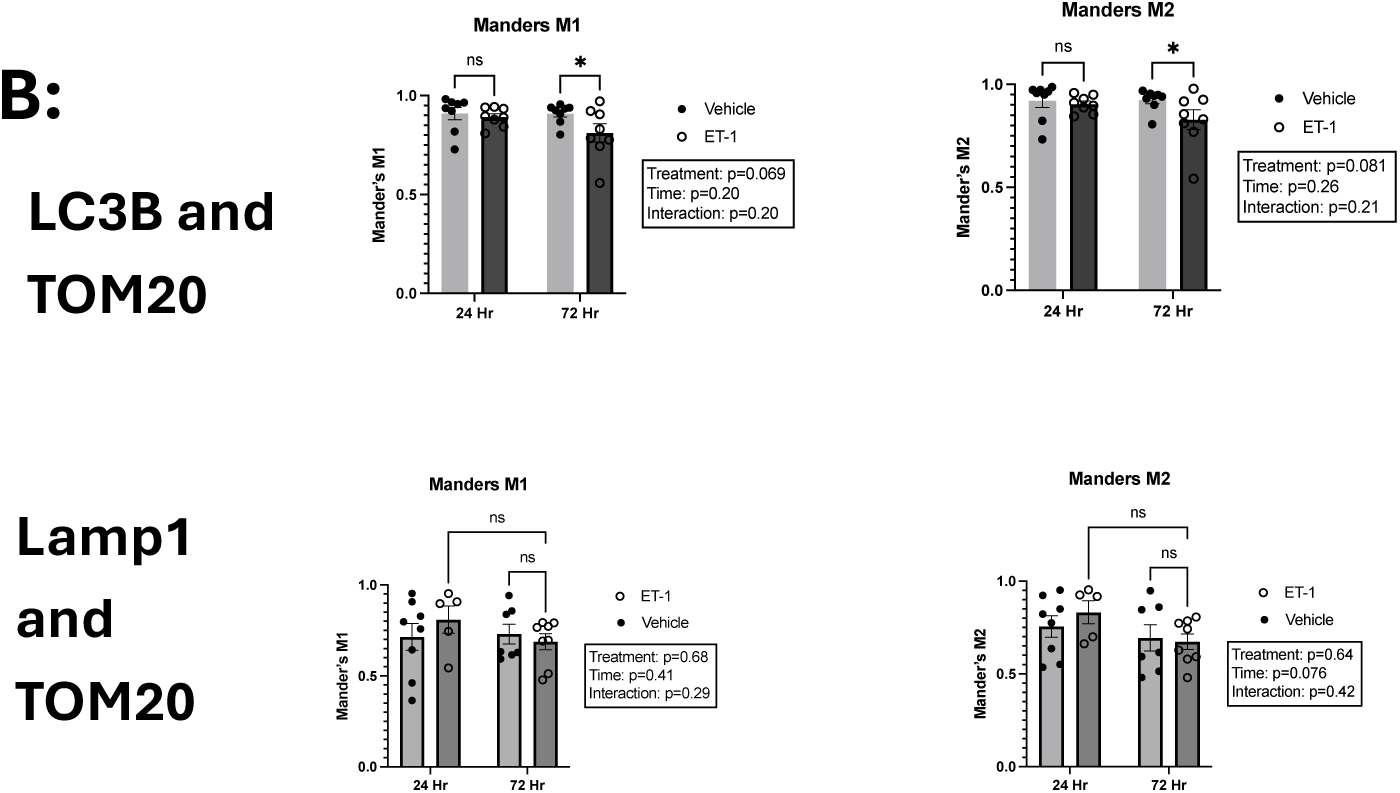
ET-1 Reduces Autophagosome Formation. Immunostaining in sagittal sections of mouse eyes injected with ET-1 or vehicle revealed that **A:** the expression of both TOM20 and LC3B were reduced 24 hours following ET-1 injection (adjusted p=0.002 and p=0.004 respectively, multiple Mann-Whiteney tests with Holm-Sidak correction). ET-1 had no effect on LAMP1 expression compared to vehicle injection. **C:** No significant differences were detected in mander’s correlation coefficient (M1 or M2) between TOM20 and LC3B or between TOM20 and LAMP1 (two-way ANOVA).

The canonical pathway of mitophagy is induced through activation of Parkin, which is phosphorylated by PINK1 stabilized on the outer mitochondrial membrane of damaged mitochondria. Parkin expression was assessed through immunohistochemistry in mice injected with ET-1 or Vehicle (Figure 4a). Additionally, an antibody targeting the activated form of Parkin (phospho-Ser63) was used to assess Parkin activation (Figure 4b). Both Parkin expression and activation was reduced 72 hours following ET-1 injection (p=0.0069 and p=0.05 respectively).

**Figure 4:**
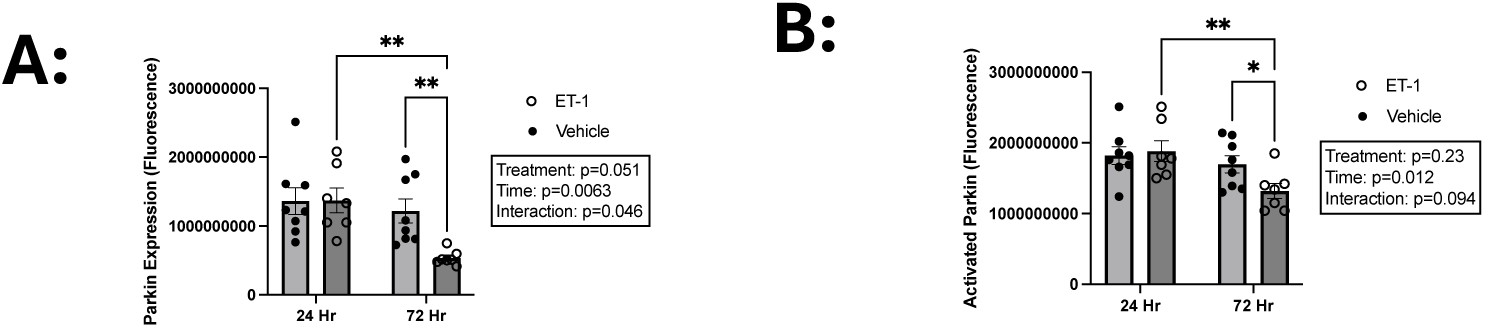
ET-1 Reduces Parkin Expression and Activation. Immunostaining in sagittal sections of mouse eyes injected with ET-1 or vehicle revealed that Parkin expression **(A)** and activation **(B)** was reduced 72 hours following ET-1 injection.

### 3.4 ET-1 Affects Mitophagy in RGCs

The MitoQC mouse was used to assess the effect of ET-1 on mitophagy. Following IVT injection of ET-1 or vehicle, one eye and optic nerve were fixed and sectioned for fluorescence microscopy. In the retina, mitophagy was assessed as mitolysosome area expressed as a percentage of the ganglion cell and nerve fiber layer area. When assessing 3 month old mice, a significant interaction effect of treatment and duration on mitolysosome percent area was detected using a two-way ANOVA (p=0.03). Post-hoc testing revealed that, when compared to vehicle treated mice, mitophagy was reduced 24 hours following ET-1 injection (p=0.01) (Figure 5a). In the optic nerve, a significant interaction effect was also detected (p=0.01). When compared with vehicle treated mice, mitophagy was increased 24 hours following ET-1 injection (p=0.003), but was reduced 72 hours following injection (p=0.04). However, injection of vehicle lead to an increase in mitophagy at 72 hours compared to 24 hours (p=0.001), limiting the ability to interpret findings at 72 hours (Figure 5b). Since glaucoma is an age-related disease, the study was repeated in older mice (>1 year old). No significant effects between treatment groups or across time points were observed in the old mice (Supplemental figure 2). Additionally, imaging of naïve animals revealed no significant age-related changes in mitophagy (Supplemental figure 2). To further explore mitophagy dynamics in response to ET-1, live imaging of retinal explants was performed in a confocal microscope. Mitolysosome area and mitolysosome number were compared across treatment groups, no significant differences between vehicle and ET-1 injected animals were detected, although there were significant trends over time (Supplemental figure 3). Further visualization and linear modeling of the live imaging data was done using R, high variability within this dataset limits the ability to interpret the data (Supplemental figure 4).

**Figure 5:**
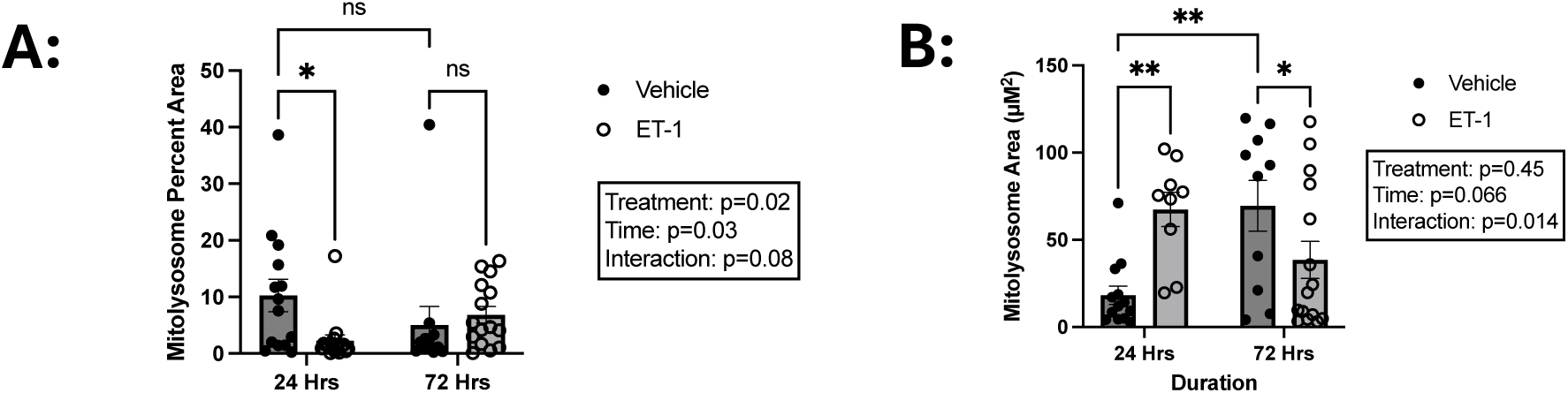
Mitophagy Changes in RGCs Following ET-1 Injection. **A:** Mitophagy was reduced in the retina 24 hours following ET-1 injection (p=0.01, two-way ANOVA with Fisher’s LSD). **B:** When compared to vehicle injected animals, mitophagy was elevated in the optic nerve 24 hours following ET-1 injection, but was reduced 72 hours following injection (p=0.003 and p=0.04 respectively, two-way ANOVA with Fisher’s LSD). However, mitophagy significantly increased over time following injection of vehicle (p=0.001).

### 3.5 IOP Affects Mitophagy in RGCs

To assess the effect of elevated intraocular pressure on mitophagy, MitoQC mice underwent intracameral injection of magnetic microbeads to clog the trabecular meshwork in one eye, the other eye served as a contralateral control. IOP was monitored, and following 1 week of IOP elevation (>16 mmHg days), eyes and optic nerves were fixed and imaged (Figure 6a). Mitophagy was elevated in the retina after IOP elevation (p=0.04) (Figure 6b). In the optic nerve, there was a decreasing trend in mitophagy following IOP elevation, but it was not significant (p=0.1) (Figure 6c).

**Figure 6:**
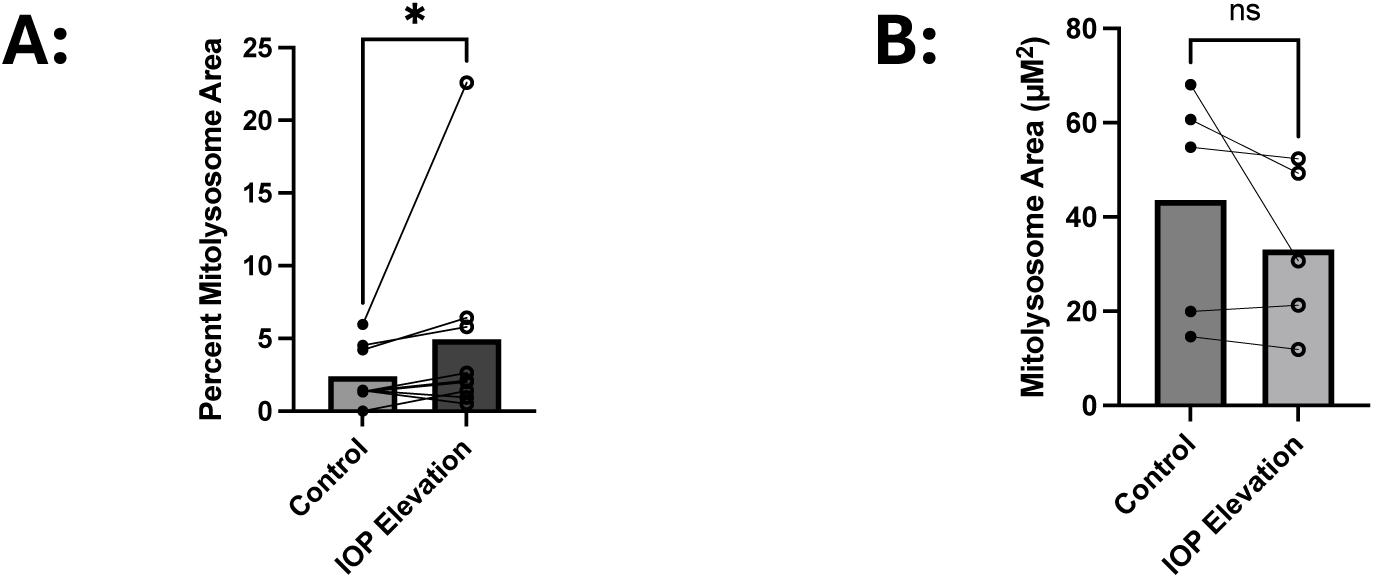
Mitophagy Changes Following IOP Elevation. **A:** In the RGC cell bodies, mitophagy was significantly increased following IOP elevation (p=0.04, Wilcoxon paired signed rank test). **B:** In the optic nerve, there was a trend towards mitophagy reductions with IOP elevation, but it was not significant (p=0.1, Wilcoxon paired signed rank test).

## Discussion

In this study, RGC loss was first seen 72 hours following ET-1 injection (Figure 1). Although the short half-life means that the initial bolus of ET-1 is long gone, it initiates a cascade that eventually leads to cell death. ET-1 has been shown to induce apoptotic cell death in rats (46). With ET-1 injection, cell death was most prominent in the central retina. This differs from the pattern of RGC loss associated with glaucoma, which sees cell loss in the peripheral retina first (45,99). The typical cell death pattern could be influenced by mechanical strain on optic nerve axons caused by increased intraocular pressure (36). The pattern of cell death in the retina is also variable, some researchers report similar RGC loss to what we see here across the retina with IOP elevation, and our previous research has found ET-1 mediated cell loss that is more pronounced in the peripheral retina (100,101).

Transmission electron microscopy of the optic nerve was used to study structural changes in mitochondria in response to ET-1. Our lab previously found ET-1 alters mitochondrial function and gene expression (51,52). As the surface area of the inner mitochondrial membrane is directly tied to the mitochondria’s capacity for oxidative phosphorylation, assessment of the cristae provides another avenue to investigate the mitochondrial dynamics of ET-1 mediated neurodegeneration (93–95). Beyond an ordinal rating scale for cristae structure, other methods for analyzing electron microscopy images of mitochondria include: surface area to volume ratio, cross sectional area, cristae density, and cristae ultrastructure (63,102). Analysis of cristae structure in optic nerve axons is confounded by the fact that mitochondria are moving through a narrow tube that is being imaged in a cross section. This leads to mitochondria appearing smaller, as the mitochondria are not being sectioned lengthwise, when cristae density and cross sectional area are typically measured in the longest dimension (63,102). Additionally, only one slice of each nerve was analyzed. The risk of slicing through an area of a mitochondria with no apparent cristae is mitigated through sampling volume. In both the vehicle and ET-1 treated animals, mitochondrial heath score tended to be lower 24 hours after injection, and tended to increase over time, but that trend was only significant in ET-1 treated animals (Figure 2b). Average mitochondrial health scores tightly clustered, inhibiting our ability to detect differences. To ensure a sufficient sampling of mitochondria to score, images were taken in areas of high axon density. This precludes us from quantitatively assessing axon death, myelin degradation, and gliosis. Our lab has previously shown axon death following ET-1 injection (47,101). Qualitatively, glial scarring and degenerated axons were observed following ET-1 injection, and became worse over time (Figure 2).

There was a marked decrease in the number of mitochondria in the optic nerve axons 24 hours following ET-1 injection (Figure 2a). ET-1 is known to impair anterograde transport of mitochondria in RGC axons, but this impairment has previously only been seen at later time points after injection (44). The decrease in mitochondrial content occurs simultaneously with an increase in mitophagy in the optic nerve (Figure 5b). The current understanding of mitophagy in RGCs is that mitophagy does not happen within the myelinated portion of the RGC axons (103). Damaged mitochondria are either degraded through mitophagy at the distal end of the axon near the synapse, sequestered into autophagosomes that undergo retrograde transport to the cell body to fuse with lysosomes, or ejected from the axon and taken up by resident glial cells and degraded in a process coined transmitophagy (103–105). In sagittal sections of optic nerves, unlike cross sections, axons and glial cells cannot be distinguished, it can be assumed that any mitophagy measured in the optic nerve sections represent the action of astrocytes and other glial cells. The drop in mitochondrial content coinciding with increased mitophagy provides evidence the ET-1 activates transmitophagy in the optic nerve. Whether this is a direct effect of ET-1, or a response to RGC distress remains to be investigated. ET-1 activates astrocytes through ET_B_, but whether IVT-injected ET-1 would reach the optic nerve is unknown (32,35,39). 72 hours following injection, optic nerves from mice injected with ET-1 had less mitophagy compared with vehicle-injected mice. It was also lower compared to ET-1 injected mice at 24 hours, although that was not significant.

Mitophagy in the optic nerve was increased in the vehicle group 72 hours after injection compared to 24 hours after injection (Figure 5b). Vehicle injection did not produce any cell death, and most factors in the retina did not change over time in vehicle injected animals. However, vehicle injection produced effects over time in the optic nerve. Transmission electron microscopy revealed that mitochondrial health drops 24 hours following vehicle injection, and improves over time. Although this was not significant in the post-hoc tests, the significant time effect in the two-way ANOVA indicates that ET-1 and vehicle treated animals follow the same general trend over time. Mitophagy in the vehicle treated animals had the opposite trend as ET-1 injected animals, with increased mitophagy at 72 hours compared with 24 hours. IVT injection, no matter the substance, can induce changes in the eye. The healing process from injection can induce changes, and the act of injection can create transient IOP increases. The mechanism behind vehicle-induced changes in the optic nerve head remain unclear.

By far the most common method for studying mitophagy is western blot assessment of the expression of proteins involved in autophagy or mitophagy, including LAMP1, ATG8, PINK1, and Parkin (79,106). However, this approach comes with several limitations. Expression of these proteins, especially Parkin and LAMP1, does not necessarily guarantee that mitophagy increases. Also, analyzing protein expression in RGCs using western blots is further complicated by the need to isolate retinal ganglion cells, as they are a small fraction of the total cells in the retina, therefore western blots of retina lysates are not necessarily indicative of protein expression in RGCs. Immunohistochemistry allows the analysis of protein expression in RGCs directly, and the localization of the protein within the cell can be analyzed as well.

In this study, the expression of TOM20, LC3B, and LAMP1 were assessed through immunohistochemistry, and were a proxy measure for the number of mitochondria, autophagosomes, and lysosomes respectively (Figure 3). The colocalization of TOM20 and LC3B was used to assess mitophagosome formation, and the colocalization of TOM20 and LAMP1 was used to assess mitolysosome formation. This method provides more information on mitophagy than simply looking at the expression of these proteins (79). We previously showed ET-1 treatment for 24 hours decreased colocalization of lysotracker and mitotracker in cultured RGCs, and TOM20 and LC3 colocalization was decreased 24 hours following IVT injection in mice, and 2 weeks of IOP elevation in rats (52). In this study we did not find significant differences in colocalization between TOM20 and LC3B or TOM20 and LAMP1. There is an increasing trend in the colocalization between TOM20 and LC3B 72 hours after both vehicle and ET-1 injection, but the ANOVA did not reach significance. Analysis of colocalization of immunofluorescence always comes with several caveats. The previous study used Mander’s Overlap Coefficient (MOC), but in this study we chose to use Mander’s Correlation Coefficient (MCC) (M1 and M2) (52). MOC is a quantifies the co-occurrence of signals regardless of the proportionality, whereas MCC is a directional measure (M1 and M2) that can be described as the proportion of factor 1 that overlaps with factor 2 (107). MOC is more often a more useful measure than Pearson’s Correlation Coefficient, as it only indicates the presence of both factors instead of their relative fluorescence intensities (107). The directional nature of MCC provides extra useful information, for factors X and Y, M1 represents the proportion of X that coincides with Y, and M2 represents the proportion of Y that coincides with X, this makes the measurement ideal when two factors may be interacting disproportionately, such as factor Y having relatively low expression, all of factor Y may be interacting factor X, but if the proteins form a heterodimer, only a fraction of factor X would be interacting with factor Y (107). In that example, M1 would show high correlation while M2 and MOC would show lower correlation as a large portion of factor Y exists in space where there is no factor X present. For the TOM20 and LC3B staining, M1 and M2 values, especially at 24 hours, are nearly identical, suggesting the above phenomenon is not a factor in the different results seen in this study. MCC is particularly sensitive to non-specific background fluorescence, necessitating the use of a background subtraction algorithm before measuring colocalization (107). In this study, as well as the previous one, a background subtraction with a rolling ball radius of 50 pixels was performed prior to colocalization analysis. It is unclear at this time the cause of the differing results.

The MitoQC mouse, along with other genetic mitophagy indicator mouse models, presents a vast improvement in the ability to detect mitophagy (79,89,106,108). A reduction in mitophagy was found 24 hours following ET-1 injection, and this reduction in mitophagy occurs at the same time as a reduction in LC3B, an autophagosome marker. This reduction in mitophagy also aligns with previous research from our group seeing declines in autophagosome formation (52). This provides further evidence that ET-1 impairs mitophagy in RGC somas, at least at early time points (Figure 5a). Interestingly, Parkin expression and activation were not changed 24 hours following ET-1 injection, suggesting that changes in mitophagy may be mediated through a parkin-independent mechanism (Figure 4). Parkin expression and activation are reduced 72 hours following ET-1 injection, when no changes in mitophagy are seen. This points to parkin-independent mitophagy, the less studied pathways, as an important regulator of RGC health (71). It is also possible that a further decline in mitophagy could occurs after longer durations (beyond 72 hours).

When examining mitophagy in mice with experimentally-induced ocular hypertension, we found mitophagy was significantly increased in the retina of IOP-elevated eyes compared to contralateral controls, but there was a decreasing trend in mitophagy in the optic nerve following IOP elevation that was not significant (Figure 6). Within eye research, it is generally accepted that the eyes operate independently and a contralateral eye can serve as a control, and this approach can be advantageous as it eliminates inter-animal variability as a confounding factor and allows the use of paired analyses (45). However, eyes are not completely independent, and experimental intervention in one eye can affect the contralateral eye (109,110). In the retina, both the IOP elevated and contralateral control eyes show lower average mitophagy than naïve or vehicle injected mice. In the optic nerve, both IOP elevated and contralateral control eyes show higher average mitophagy than naïve or vehicle injected mice, except for 72 hours following vehicle injection in young mice. This could indicate that mitophagy decreases in the retina and increases in the optic nerve with IOP elevation, and the contralateral eye experiences similar effects. This could also explain the small effect size seen when comparing mitophagy in IOP elevated eyes with contralateral controls.

Another factor that limits interpretation of mitophagy with IOP elevation in this study is the high variability within the sample and relatively low sample size. Some of the variability could be due to differential IOP exposure. In this study, mice were IOP elevated for 1 week. After tissue collection, IOP exposure was calculated as the area under the curve of the IOP increase (mmHg*Days), and a cutoff of 16 mmHg days was chosen for inclusion in the study, representing a >2 mmHg IOP elevation for 7 days. While most mice had an IOP exposure of 19-23 mmHg*Days, a couple animals had higher IOP exposure. The mitophagy changes could be proportional to the IOP exposure, but we are not significantly powered to test that hypothesis.

Our study examined mitophagy at a relatively shorter time frame of IOP elevation, and a much smaller IOP exposure. The goal was to examine mitophagy early in the disease progression, before cell loss occurs. This may explain why we find mitophagy increasing with IOP elevation when others report reductions in mitophagy after IOP elevation sustained for multiple weeks (88). Ma et al. (2025) found results similar to ours, finding markers of mitophagy initially increasing after induction of IOP, but tapering off and eventually becoming reduced compared to controls after 2 weeks of IOP elevation (87). Our goal in examining this early time point with mild IOP elevation was to see if similar perturbations of the mitophagy pathway are seen at early time points following IOP elevation as seen 24 hours following ET-1 injection. These insults were hypothesized to produce similar results since both end points occur before significant RGC loss occurs, this would produce insight into the dysfunctional state that eventually causes RGC death. The different results seen with IOP elevation and ET-1 injection could be due to each insult having a different pattern and timeframe of damage. ET-1 injection significantly accelerates the time frame for RGC loss, so perhaps the insult is more harsh than the mild IOP elevation we tested. Another explanation could simply be the heterogeneous nature of glaucoma. While ET-1 is increased in glaucoma, and we show here that ET-1 affects mitophagy, other factors are affected by increased IOP and could affect mitophagy in other ways. In the optic nerve, mitophagy was increased 24 hours following ET-1 injection, but there was a trend towards mitophagy reductions following IOP elevation. This difference could be due to the different pathway of damage in the optic nerve. Increased IOP puts mechanical strain on the optic nerve head, which is absent with ET-1 administration alone, this represents a separate insult which cells respond to, and could lead to different responses compared to ET-1 alone, even though they both lead to fibrotic remodeling of the optic nerve head region (36,41).

The mechanism for ET-1 mediated mitophagy changes in RGCs has not been fully elucidated. ET_B_ selective agonists produce mitochondrial dysfunction and cell death (45,53,111,112). However, overexpression of ET_A_ also produces RGC death, and activation of one endothelin receptor increases the expression of the other receptor (45,49). In addition to IP_3_-mediated calcium release, ET_B_ signaling activates a variety of signaling cascades and transcription factors through G_0_, G_i_, and G_q_ proteins (113,114). ET_B_-mediated c-Jun N-terminal kinase 1 (JNK1) activation and subsequent phosphorylation of c-Jun has been shown in a variety of cell types, including astrocytes (42,114–116). JNK/c-Jun activation also occurs in response to ET-1 in human non-pigmented ciliary body epithelial cells, and is responsible for the upregulation of both ET receptors in response to receptor activation (117). c-Jun phosphorylation was increased following ET-1 and ET-3 administration in cultured RGCs (50). Furthermore, JNK2 knockout in mice protected RGCs and prevented increased phosphorylation of c-Jun following IVT injection of ET-1 (101). In addition to c-Jun, JNK1 and JNK2 can phosphorylate BCL2/adenovirus e1B 19 kDa protein interacting protein 3 (BNIP3), stabilizing it on the outer mitochondrial membrane where it directly interacts with LC3B initiating Parkin-independent mitophagy (118,119). This pathway could be responsible for the disconnect between observed trends in mitophagy and Parkin expression and activation. Mitophagy was reduced 24 hours following ET-1 injection, even though there was no change in Parkin expression or activation. However, mitophagy did not differ between ET-1 and vehicle injected animals 72 hours following injection, despite a marked reduction in Parkin expression and activation with ET-1 treatment at that timepoint.

Since age is a risk factor for the development of primary open angle glaucoma, we sought to evaluate the effect of ET-1 on mitophagy in aged mice (3). Mitophagy impairments have been seen in aging and implicated in the pathogenesis of many age-related diseases including neurodegenerative diseases (77,79,120,121). Older mice (>1 year old), showed no changes in mitophagy in response to ET-1 injection, nor showed changing mitophagy across time points. This was true for the optic nerve and the retina. In naïve animals, there was no difference in mitophagy between young and old mice. It is possible that older mice had a reduced capacity to change mitophagy, no matter the direction of the change.

Live imaging of MitoQC mouse retinas following ET-1 injection was conducted to monitor mitophagic flux in real time. The high magnification combined with a large sampling of individual retinal ganglion cells and Z-stacking to quantify the entire cell was predicted to provide a clearer look at mitophagy in retinal ganglion cells. Mitolysosome number and mitolysosome area was examined in retinas 24, 48, or 72 hours following ET-1 injection in young MitoQC mice, and no differences were seen. Additionally, older mice were examined 24 and 72 hours following ET-1 injection, and no differences were seen. Naïve animals (young and old) were compared and no difference was seen. This live imaging data had high variability, even when examining individual cells within the same animal, there was a wide range in mitophagy levels. Additionally, this variability was not normally distributed. It is possible that this variability is indicative of subtypes of RGCs with different basal mitophagy levels, several subtypes of RGCs have been identified to date and these subtypes may respond differently to cellular stress (122,123). Additionally, in many samples green fluorescence intensity far exceeded red fluorescence activity, and green fluorescence seemed to exist in areas without red fluorescence. This phenomenon was not seen when examining sagittal sections. Autofluorescense is common in the retinal pigment epithelium and photoreceptors, although the pinhole aperture of the confocal microscope should limit spurious fluorescence from outside the focal plane. Thresholding was used to identify all pixels with red fluorescence, and mitophagy was quantified only in those pixels, although if this green autofluorescence happened to occur in the same pixel as red fluorescence, the mitophagosome count may appear lower than it actually is. Future live imaging of mitophagy using MitoQC mice would best be conducted in isolated, cultured, RGCs as opposed to retinal explants.

## Conclusion

ET-1 injection into the retina causes a signaling cascade in RGCs that ultimately leads to mitochondrial dysfunction and cell death. The data presented here show that mitophagy in the RGC cell bodies and mitophagy in the optic nerve, presumably in glial cells, follow different trends. With ET-1, mitochondrial health initially decreases in RGC axons, but mitochondrial health recovers over time in surviving cells. This, combined with the increased mitophagy throughout the nerve and decreased mitochondria numbers in the axons suggests that transmitophagy is the mechanism responsible for that recovery. Since mitophagy was not increased in the optic nerve following IOP elevation, increasing transmitophagy may be a viable strategy for preventing axon loss during IOP elevation.

Meanwhile, in the RGC cell bodies, mitophagy was increased following mild IOP elevation, and could represent an initial period of compensation for the IOP-induced cell stress, which eventually devolves. Mitophagy was reduced 24 hours following ET-1 injection, and although that reduction was alleviated 72 hours following injection, it could be an early contributing factor to the cell death following ET-1 treatment. ET-1 mediated changes in mitophagy were found to be independent of Parkin expression and activation, and further studies should be conducted into the role of the Parkin-independent mitophagy pathways in response to cellular stress in RGCs, as these may produce valid targets for neuroprotective treatments.

## Acknowledgements

These studies were supported by the National Institutes of Health R01EY028179 and T32AG020494. We would like to thank Chantal Allamargot (University of Iowa) and Anne-Marie Brun (UNTHSC) for assistance with experiments.

## References

1. Weinreb RN, Leung CKS, Crowston JG, Medeiros FA, Friedman DS, Wiggs JL, et al. Primary open-angle glaucoma. Nat Rev Dis Primer. 2016 Sep 22;2(1):1. doi:10.1038/nrdp.2016.67

2. Quigley HA. Neuronal death in glaucoma. Prog Retin Eye Res. 1999 Jan 1;18(1):39–57. doi:10.1016/S1350-9462(98)00014-7

3. Boland MV, Quigley HA. Risk Factors and Open-angle Glaucoma: Classification and Application. J Glaucoma. 2007 Jul;16(4):406. doi:10.1097/IJG.0b013e31806540a1

4. Lusthaus J, Goldberg I. Current management of glaucoma. Med J Aust. 2019;210(4):4. doi:10.5694/mja2.50020

5. Liu P, Wang F, Song Y, Wang M, Zhang X. Current situation and progress of drugs for reducing intraocular pressure. Ther Adv Chronic Dis. 2022 Jan;13:204062232211403. doi:10.1177/20406223221140392

6. Gutiérrez Martín LC. Update on the diagnosis and treatment of normotensive glaucoma. Arch Soc Esp Oftalmol Engl Ed. 2023 Jun;98(6):344–50. doi:10.1016/j.oftale.2023.05.004

7. Khawaja AP, Campbell JH, Kirby N, Chandwani HS, Keyzor I, Parekh M, et al. Real-World Outcomes of Selective Laser Trabeculoplasty in the United Kingdom. Ophthalmology. 2020 Jun;127(6):6. doi:10.1016/j.ophtha.2019.11.017

8. Bovell AM, Damji KF, Hodge WG, Rock WJ, Buhrmann RR, Pan YI. Long term effects on the lowering of intraocular pressure: selective laser or argon laser trabeculoplasty? Can J Ophthalmol. 2011 Oct;46(5):5. doi:10.1016/j.jcjo.2011.07.016

9. Jampel HD, Chon BH, Stamper R, Packer M, Han Y, Nguyen QH, et al. Effectiveness of Intraocular Pressure–Lowering Medication Determined by Washout. JAMA Ophthalmol. 2014 Apr 1;132(4):4. doi:10.1001/jamaophthalmol.2013.7677

10. Stallworth JY, O’Brien KS, Han Y, Oatts JT. Efficacy of Ahmed and Baerveldt glaucoma drainage device implantation in the pediatric population: A systematic review and meta-analysis. Surv Ophthalmol. 2023 Jul;68(4):4. doi:10.1016/j.survophthal.2023.01.010

11. Malek I, Sayadi J, Choura R, Mekni M, Rayhane H, Khairallah M, et al. Long-Term Results of Combined Trabeculotomy Trabeculectomy in Primary Congenital Glaucoma. J Glaucoma. 2023 Oct;32(10):10. doi:10.1097/IJG.0000000000002229

12. DuBiner HB, Mroz M, Shapiro AM, Dirks MS. A comparison of the efficacy and tolerability of brimonidine and latanoprost in adults with open-angle glaucoma or ocular hypertension: A three-month, multicenter, randomized, double-masked, parallel-group trial. Clin Ther. 2001 Dec;23(12):12. doi:10.1016/S0149-2918(01)80150-8

13. Sakaorat P, Mohamed-Noriega J, Sharara A, Daniel MC, Brookes J. Cyclodiode Laser as the First Surgical Approach in Childhood Glaucoma Under the Age of 8 Years. J Glaucoma. 2021 Apr;30(4):4. doi:10.1097/IJG.0000000000001754

14. Uva MG, Avitabile T, Reibaldi M, Bucolo C, Drago F, Quaranta L, et al. Long-term efficacy of latanoprost in primary congenital glaucoma. Eye. 2014 Jan;28(1):53–7. doi:10.1038/eye.2013.232

15. Peters D, Bengtsson B, Heijl A. Lifetime Risk of Blindness in Open-Angle Glaucoma. Am J Ophthalmol. 2013 Oct;156(4):4. doi:10.1016/j.ajo.2013.05.027

16. Susanna R, De Moraes CG, Cioffi GA, Ritch R. Why Do People (Still) Go Blind from Glaucoma? Transl Vis Sci Technol. 2015 Mar;4(2):2. doi:10.1167/tvst.4.2.1

17. Pascale A, Drago F, Govoni S. Protecting the retinal neurons from glaucoma: Lowering ocular pressure is not enough. Pharmacol Res. 2012 Jul 1;66(1):19–32. doi:10.1016/j.phrs.2012.03.002

18. Itoh Y, Yanagisawa M, Ohkubo S, Kimura C, Kosaka T, Inoue A, et al. Cloning and sequence analysis of cDNA encoding the precursor of a human endothelium-derived vasoconstrictor peptide, endothelin: identity of human and porcine endothelin. FEBS Lett. 1988 Apr 25;231(2):440–4. doi:10.1016/0014-5793(88)80867-6 PubMed PMID: 3282927.

19. Masaki T. The discovery, the present state, and the future prospects of endothelin. J Cardiovasc Pharmacol. 1989;13 Suppl 5:S1–4; discussion S18. doi:10.1097/00005344-198900135-00002 PubMed PMID: 2473280.

20. Moore R, Linas S. Endothelin antagonists and resistant hypertension in chronic kidney disease. Curr Opin Nephrol Hypertens. 2010 Sep;19(5):5. doi:10.1097/MNH.0b013e32833a7a25

21. Lazich I, Bakris GL. Endothelin Antagonism in Patients with Resistant Hypertension and Hypertension Nephropathy. Endothel Ren Physiol Dis. 2011;172:223–34. doi:10.1159/000328988 PubMed PMID: 21894002.

22. Kohan DE, Barton M. Endothelin and Endothelin Antagonists in Chronic Kidney Disease. Kidney Int. 2014 Nov;86(5):5. doi:10.1038/ki.2014.143 PubMed PMID: 24805108; PubMed Central PMCID: PMC4216619.

23. Dammanahalli KJ, Sun Z. Endothelins and Nadph Oxidases in the Cardiovascular System. Clin Exp Pharmacol Physiol. 2008;35(1):2–6. doi:10.1111/j.1440-1681.2007.04830.x

24. MacCumber MW, D’Anna SA. Endothelin Receptor-Binding Subtypes in the Human Retina and Choroid. Arch Ophthalmol. 1994 Sep 1;112(9):9. doi:10.1001/archopht.1994.01090210119024

25. Chauhan BC. Endothelin and its potential role in glaucoma. Can J Ophthalmol. 2008 Jun 1;43(3):356–60. doi:10.3129/i08-060

26. Krishnamoorthy RR, Prasanna G, Dauphin R, Hulet C, Agarwal N, Yorio T. Regulation of Na,K-ATPase Expression by Endothelin-1 in Transformed Human Ciliary Non-Pigmented Epithelial (HNPE) Cells. J Ocul Pharmacol Ther. 2003 Oct;19(5):465–81. doi:10.1089/108076803322473024

27. Zhang X, Krishnamoorthy RR, Prasanna G, Narayan S, Clark A, Yorio T. Dexamethasone regulates endothelin-1 and endothelin receptors in human non-pigmented ciliary epithelial (HNPE) cells. Exp Eye Res. 2003 Mar;76(3):261–72. doi:10.1016/S0014-4835(02)00323-8

28. Narayan S, Prasanna G, Krishnamoorthy RR, Zhang X, Yorio T. Endothelin-1 Synthesis and Secretion in Human Retinal Pigment Epithelial Cells (ARPE-19): Differential Regulation by Cholinergics and TNF-α. Invest Ophthalmol Vis Sci. 2003 Nov 1;44(11):4885–94. doi:10.1167/iovs.03-0387

29. Narayan S, Prasanna G, Tchedre K, Krishnamoorthy R, Yorio T. Thrombin-Induced Endothelin-1 Synthesis and Secretion in Retinal Pigment Epithelial Cells Is Rho Kinase Dependent. J Ocul Pharmacol Ther. 2010 Oct;26(5):389–97. doi:10.1089/jop.2010.0072 PubMed PMID: 20874501; PubMed Central PMCID: PMC2956378.

30. Hostenbach S, D’haeseleer M, Kooijman R, De Keyser J. The pathophysiological role of astrocytic endothelin-1. Prog Neurobiol. 2016 Sep;144:88–102. doi:10.1016/j.pneurobio.2016.04.009 PubMed PMID: 27132521.

31. Desai D, He S, Yorio T, Krishnamoorthy RR, Prasanna G. Hypoxia augments TNF-α-mediated endothelin-1 release and cell proliferation in human optic nerve head astrocytes. Biochem Biophys Res Commun. 2004 Jun;318(3):642–8. doi:10.1016/j.bbrc.2004.04.073

32. Prasanna G, Krishnamoorthy R, Yorio T. Endothelin, Astrocytes and Glaucoma. Exp Eye Res. 2011 Aug;93(2):170–7. doi:10.1016/j.exer.2010.09.006 PubMed PMID: 20849847; PubMed Central PMCID: PMC3046320.

33. Tezel G, Kass MA, Kolker AE, Becker B, Wax MB. Plasma and aqueous humor endothelin levels in primary open-angle glaucoma. J Glaucoma. 1997 Apr;6(2):83–9. PubMed PMID: 9098815.

34. Källberg ME, Brooks DE, Garcia-Sanchez GA, Komàromy AM, Szabo NJ, PhD LT. Endothelin 1 Levels in the Aqueous Humor of Dogs With Glaucoma. J Glaucoma. 2002 Apr;11(2):105.

35. Prasanna G, Hulet C, Desai D, Krishnamoorthy RR, Narayan S, Brun AM, et al. Effect of elevated intraocular pressure on endothelin-1 in a rat model of glaucoma. Pharmacol Res. 2005 Jan;51(1):41–50. doi:10.1016/j.phrs.2004.04.006

36. Morrison JC, Moore CG, Deppmeier LMH, Gold BG, Meshul CK, Johnson EC. A Rat Model of Chronic Pressure-induced Optic Nerve Damage. Exp Eye Res. 1997 Jan 1;64(1):85–96. doi:10.1006/exer.1996.0184

37. Li S, Zhang A, Cao W, Sun X. Elevated Plasma Endothelin-1 Levels in Normal Tension Glaucoma and Primary Open-Angle Glaucoma: A Meta-Analysis. J Ophthalmol. 2016;2016:2678017. doi:10.1155/2016/2678017 PubMed PMID: 27965889; PubMed Central PMCID: PMC5124679.

38. Choritz L, Machert M, Thieme H. Correlation of endothelin-1 concentration in aqueous humor with intraocular pressure in primary open angle and pseudoexfoliation glaucoma. Invest Ophthalmol Vis Sci. 2012 Oct 23;53(11):7336–42. doi:10.1167/iovs.12-10216 PubMed PMID: 23036995.

39. Prasanna G, Krishnamoorthy R, Clark AF, Wordinger RJ, Yorio T. Human Optic Nerve Head Astrocytes as a Target for Endothelin-1. Invest Ophthalmol Vis Sci. 2002 Aug 1;43(8):2704–13.

40. Rao VR, Krishnamoorthy RR, Yorio T. Endothelin-1, endothelin A and B receptor expression and their pharmacological properties in GFAP negative human lamina cribrosa cells. Exp Eye Res. 2007 Jun;84(6):1115–24. doi:10.1016/j.exer.2007.02.010

41. Rao VR, Krishnamoorthy RR, Yorio T. Endothelin-1 Mediated Regulation of Extracellular Matrix Collagens in Cells of Human Lamina Cribrosa. Exp Eye Res. 2008 Jun;86(6):886–94. doi:10.1016/j.exer.2008.03.003 PubMed PMID: 18420197; PubMed Central PMCID: PMC2467437.

42. Gadea A, Schinelli S, Gallo V. Endothelin-1 Regulates Astrocyte Proliferation and Reactive Gliosis via a JNK/c-Jun Signaling Pathway. J Neurosci. 2008 Mar 5;28(10):2394– 408. doi:10.1523/JNEUROSCI.5652-07.2008 PubMed PMID: 18322086; PubMed Central PMCID: PMC2695974.

43. Hammond TR, McEllin B, Morton PD, Raymond M, Dupree J, Gallo V. Endothelin-B Receptor Activation in Astrocytes Regulates the Rate of Oligodendrocyte Regeneration during Remyelination. Cell Rep. 2015 Dec 15;13(10):2090–7. doi:10.1016/j.celrep.2015.11.002

44. Stokely ME, Yorio T, King MA. Endothelin-1 modulates anterograde fast axonal transport in the central nervous system. J Neurosci Res. 2005;79(5):5. doi:10.1002/jnr.20383

45. Minton AZ, Phatak NR, Stankowska DL, He S, Ma HY, Mueller BH, et al. Endothelin B Receptors Contribute to Retinal Ganglion Cell Loss in a Rat Model of Glaucoma. PLoS ONE. 2012 Aug 20;7(8):e43199. doi:10.1371/journal.pone.0043199 PubMed PMID: 22916224; PubMed Central PMCID: PMC3423444.

46. Lau J, Dang M, Hockmann K, Ball AK. Effects of acute delivery of endothelin-1 on retinal ganglion cell loss in the rat. Exp Eye Res. 2006 Jan 1;82(1):132–45. doi:10.1016/j.exer.2005.06.002

47. Kodati B, Zhang W, He S, Pham JH, Beall KJ, Swanger ZE, et al. The endothelin receptor antagonist macitentan ameliorates endothelin-mediated vasoconstriction and promotes the survival of retinal ganglion cells in rats. Front Ophthalmol [Internet]. 2023 [cited 2024 Mar 1];3. Available from: https://www.frontiersin.org/articles/10.3389/fopht.2023.1185755

48. Kodati B, McGrady NR, Jefferies HB, Stankowska DL, Krishnamoorthy RR. Oral administration of a dual ETA/ETB receptor antagonist promotes neuroprotection in a rodent model of glaucoma. Mol Vis. 2022 Aug 7;28:165–77. PubMed PMID: 36274816; PubMed Central PMCID: PMC9491150.

49. McGrady NR, Minton AZ, Stankowska DL, He S, Jefferies HB, Krishnamoorthy RR. Upregulation of the endothelin A (ETA) receptor and its association with neurodegeneration in a rodent model of glaucoma. BMC Neurosci. 2017 Mar 1;18:27. doi:10.1186/s12868-017-0346-3 PubMed PMID: 28249604; PubMed Central PMCID: PMC5333388.

50. He S, Park YH, Yorio T, Krishnamoorthy RR. Endothelin-Mediated Changes in Gene Expression in Isolated Purified Rat Retinal Ganglion Cells. Invest Ophthalmol Vis Sci. 2015 Sep;56(10):6144–61. doi:10.1167/iovs.15-16569 PubMed PMID: 26397462; PubMed Central PMCID: PMC5102497.

51. Chaphalkar RM, Stankowska DL, He S, Kodati B, Phillips N, Prah J, et al. Endothelin-1 Mediated Decrease in Mitochondrial Gene Expression and Bioenergetics Contribute to Neurodegeneration of Retinal Ganglion Cells. Sci Rep. 2020 Feb 27;10:3571. doi:10.1038/s41598-020-60558-6 PubMed PMID: 32107448; PubMed Central PMCID: PMC7046667.

52. Chaphalkar RM, Kodati B, Maddineni P, He S, Brooks CD, Stankowska DL, et al. A Reduction in Mitophagy Is Associated with Glaucomatous Neurodegeneration in Rodent Models of Glaucoma. Int J Mol Sci. 2024 Dec 4;25(23):13040. doi:10.3390/ijms252313040 PubMed PMID: 39684751; PubMed Central PMCID: PMC11642561.

53. Tam SW, Feng R, Lau WKW, Law ACK, Yeung PKK, Chung SK. Endothelin type B receptor promotes cofilin rod formation and dendritic loss in neurons by inducing oxidative stress and cofilin activation. J Biol Chem. 2019 Aug 16;294(33):33. doi:10.1074/jbc.RA118.005155

54. Schrier SA, Falk MJ. Mitochondrial disorders and the eye. Curr Opin Ophthalmol. 2011 Sep;22(5):325–31. doi:10.1097/ICU.0b013e328349419d PubMed PMID: 21730846; PubMed Central PMCID: PMC3652603.

55. Klemmensen MM, Borrowman SH, Pearce C, Pyles B, Chandra B. Mitochondrial dysfunction in neurodegenerative disorders. Neurotherapeutics. 2024 Jan 1;21(1):e00292. doi:10.1016/j.neurot.2023.10.002

56. Terluk MR, Kapphahn RJ, Soukup LM, Gong H, Gallardo C, Montezuma SR, et al. Investigating mitochondria as a target for treating age-related macular degeneration. J Neurosci Off J Soc Neurosci. 2015 May 6;35(18):7304–11. doi:10.1523/JNEUROSCI.0190-15.2015 PubMed PMID: 25948278; PubMed Central PMCID: PMC4420790.

57. Karunadharma PP, Nordgaard CL, Olsen TW, Ferrington DA. Mitochondrial DNA damage as a potential mechanism for age-related macular degeneration. Invest Ophthalmol Vis Sci. 2010 Nov;51(11):5470–9. doi:10.1167/iovs.10-5429 PubMed PMID: 20505194; PubMed Central PMCID: PMC3061495.

58. Hollyfield JG, Bonilha VL, Rayborn ME, Yang X, Shadrach KG, Lu L, et al. Oxidative damage-induced inflammation initiates age-related macular degeneration. Nat Med. 2008 Feb;14(2):194–8. doi:10.1038/nm1709 PubMed PMID: 18223656; PubMed Central PMCID: PMC2748836.

59. Brown EE, DeWeerd AJ, Ildefonso CJ, Lewin AS, Ash JD. Mitochondrial oxidative stress in the retinal pigment epithelium (RPE) led to metabolic dysfunction in both the RPE and retinal photoreceptors. Redox Biol. 2019 Jun;24:101201. doi:10.1016/j.redox.2019.101201 PubMed PMID: 31039480; PubMed Central PMCID: PMC6488819.

60. Osborne NN, del Olmo-Aguado S. Maintenance of retinal ganglion cell mitochondrial functions as a neuroprotective strategy in glaucoma. Curr Opin Pharmacol. 2013 Feb 1;Neurosciences13(1):1. doi:10.1016/j.coph.2012.09.002

61. Inman DM, Harun-Or-Rashid M. Metabolic Vulnerability in the Neurodegenerative Disease Glaucoma. Front Neurosci. 2017;11:146. doi:10.3389/fnins.2017.00146 PubMed PMID: 28424571; PubMed Central PMCID: PMC5371671.

62. Yang S, Park JH, Lu HC. Axonal energy metabolism, and the effects in aging and neurodegenerative diseases. Mol Neurodegener. 2023 Jul 20;18(1):49. doi:10.1186/s13024-023-00634-3 PubMed PMID: 37475056; PubMed Central PMCID: PMC10357692.

63. Ju WK, Perkins GA, Kim KY, Bastola T, Choi WY, Choi SH. Glaucomatous optic neuropathy: Mitochondrial dynamics, dysfunction and protection in retinal ganglion cells. Prog Retin Eye Res. 2023 Jul;95:101136. doi:10.1016/j.preteyeres.2022.101136

64. Zhang ZQ, Xie Z, Chen SY, Zhang X. Mitochondrial dysfunction in glaucomatous degeneration. Int J Ophthalmol. 2023 May 18;16(5):811–23. doi:10.18240/ijo.2023.05.20 PubMed PMID: 37206187; PubMed Central PMCID: PMC10172101.

65. Pinazo-Durán MD, Shoaie-Nia K, Zanón-Moreno V, Sanz-González SM, Benítez del Castillo J, García-Medina JJ. Strategies to Reduce Oxidative Stress in Glaucoma Patients. Curr Neuropharmacol. 2018 Aug;16(7):7. doi:10.2174/1570159X15666170705101910 PubMed PMID: 28677495; PubMed Central PMCID: PMC6120109.

66. Osborne NN, Núñez-Álvarez C, Joglar B, del Olmo-Aguado S. Glaucoma: Focus on mitochondria in relation to pathogenesis and neuroprotection. Eur J Pharmacol. 2016 Sep;787:127–33. doi:10.1016/j.ejphar.2016.04.032

67. Ryan AK, Rich W, Reilly MA. Oxidative stress in the brain and retina after traumatic injury. Front Neurosci. 2023;17:1021152. doi:10.3389/fnins.2023.1021152 PubMed PMID: 36816125; PubMed Central PMCID: PMC9935939.

68. Tribble JR, Otmani A, Sun S, Ellis SA, Cimaglia G, Vohra R, et al. Nicotinamide provides neuroprotection in glaucoma by protecting against mitochondrial and metabolic dysfunction. Redox Biol. 2021 Jul;43:101988. doi:10.1016/j.redox.2021.101988

69. Williams PA, Harder JM, Foxworth NE, Cochran KE, Philip VM, Porciatti V, et al. Vitamin B _3_ modulates mitochondrial vulnerability and prevents glaucoma in aged mice. Science. 2017 Feb 17;355(6326):756–60. doi:10.1126/science.aal0092

70. Jassim AH, Inman DM, Mitchell CH. Crosstalk Between Dysfunctional Mitochondria and Inflammation in Glaucomatous Neurodegeneration. Front Pharmacol. 2021;12:699623. doi:10.3389/fphar.2021.699623 PubMed PMID: 34366851; PubMed Central PMCID: PMC8334009.

71. Villa E, Marchetti S, Ricci JE. No Parkin Zone: Mitophagy without Parkin. Trends Cell Biol. 2018 Nov 1;28(11):11. doi:10.1016/j.tcb.2018.07.004

72. Ryan TA, Tumbarello DA. Optineurin: A Coordinator of Membrane-Associated Cargo Trafficking and Autophagy. Front Immunol. 2018 May 15;9:1024. doi:10.3389/fimmu.2018.01024 PubMed PMID: 29867991; PubMed Central PMCID: PMC5962687.

73. Wong YC, Holzbaur ELF. Temporal dynamics of PARK2/parkin and OPTN/optineurin recruitment during the mitophagy of damaged mitochondria. Autophagy. 2015 Mar 24;11(2):422–4. doi:10.1080/15548627.2015.1009792 PubMed PMID: 25801386; PubMed Central PMCID: PMC4502688.

74. Ashrafi G, Schwarz TL. The pathways of mitophagy for quality control and clearance of mitochondria. Cell Death Difffer. 2013 Jan;20(1):1. doi:10.1038/cdd.2012.81

75. Popov LD. Mitochondrial biogenesis: An update. J Cell Mol Med. 2020 May;24(9):4892–9. doi:10.1111/jcmm.15194 PubMed PMID: 32279443; PubMed Central PMCID: PMC7205802.

76. Zamponi N, Zamponi E, Cannas SA, Billoni OV, Helguera PR, Chialvo DR. Mitochondrial network complexity emerges from fission/fusion dynamics. Sci Rep. 2018 Jan 10;8(1):363. doi:10.1038/s41598-017-18351-5 PubMed PMID: 29321534; PubMed Central PMCID: PMC5762699.

77. Wang Y, Liu N, Lu B. Mechanisms and roles of mitophagy in neurodegenerative diseases. CNS Neurosci Ther. 2019;25(7):7. doi:10.1111/cns.13140

78. Quinn PMJ, Moreira PI, Ambrósio AF, Alves CH. PINK1/PARKIN signalling in neurodegeneration and neuroinflammation. Acta Neuropathol Commun. 2020 Dec;8(1):1. doi:10.1186/s40478-020-01062-w

79. Brooks CD, Kodati B, Stankowska DL, Krishnamoorthy RR. Role of mitophagy in ocular neurodegeneration. Front Neurosci. 2023 Oct 27;17. doi:10.3389/fnins.2023.1299552

80. Alka K, Kumar J, Kowluru RA. Impaired mitochondrial dynamics and removal of the damaged mitochondria in diabetic retinopathy. Front Endocrinol. 2023;14:1160155. doi:10.3389/fendo.2023.1160155 PubMed PMID: 37415667; PubMed Central PMCID: PMC10320727.

81. Zhou P, Xie W, Meng X, Zhai Y, Dong X, Zhang X, et al. Notoginsenoside R1 Ameliorates Diabetic Retinopathy through PINK1-Dependent Activation of Mitophagy. Cells. 2019 Mar 2;8(3):213. doi:10.3390/cells8030213 PubMed PMID: 30832367; PubMed Central PMCID: PMC6468581.

82. Hombrebueno JR, Cairns L, Dutton LR, Lyons TJ, Brazil DP, Moynagh P, et al. Uncoupled turnover disrupts mitochondrial quality control in diabetic retinopathy. JCI Insight. 2019 Dec 5;4(23):e129760, 129760. doi:10.1172/jci.insight.129760 PubMed PMID: 31661466; PubMed Central PMCID: PMC6962019.

83. Obanina NA, Bgatova NP, Eremina AV, Trunov AN, Chernykh VV. Autophagy in Human Retinal Neurons in Glaucoma. Bull Exp Biol Med. 2022 Aug;173(4):4. doi:10.1007/s10517-022-05563-7

84. Zeng W, Wang W, Wu S, Zhu X, Zheng T, Chen X, et al. Mitochondria and Autophagy Dysfunction in Glucocorticoid-Induced Ocular Hypertension/Glaucoma Mice Model. Curr Eye Res. 2020 Feb 1;45(2):2. doi:10.1080/02713683.2019.1657462

85. Hass DT, Barnstable CJ. Mitochondrial Uncoupling Protein 2 Knock-out Promotes Mitophagy to Decrease Retinal Ganglion Cell Death in a Mouse Model of Glaucoma. J Neurosci. 2019 May 1;39(18):3582–96. doi:10.1523/JNEUROSCI.2702-18.2019 PubMed PMID: 30814312; PubMed Central PMCID: PMC6495138.

86. Sridevi Gurubaran I, Viiri J, Koskela A, Hyttinen JMT, Paterno JJ, Kis G, et al. Mitophagy in the Retinal Pigment Epithelium of Dry Age-Related Macular Degeneration Investigated in the NFE2L2/PGC-1α-/- Mouse Model. Int J Mol Sci. 2020 Mar 13;21(6):1976. doi:10.3390/ijms21061976 PubMed PMID: 32183173; PubMed Central PMCID: PMC7139489.

87. Ma H, Hu X, Zhang J, Lian W, Wang D, Zhang L, et al. Fucoxanthin protects retinal ganglion cells and regulates Parkin-mediated mitophagy in experimental glaucoma. BMJ Open Ophthalmol. 2025 Aug;10(1):e002126. doi:10.1136/bmjophth-2024-002126

88. Maddineni P, Kaipa BR, Kodati B, Kesavan K, Li L, Millar JC, et al. Pharmacological restoration of impaired autophagy in retinal ganglion cells prevents abnormal mitochondrial accumulation and glaucomatous neurodegeneration. Mol Neurodegener. 2026 May 16. doi:10.1186/s13024-026-00950-4

89. McWilliams TG, Prescott AR, Allen GFG, Tamjar J, Munson MJ, Thomson C, et al. mito-QC illuminates mitophagy and mitochondrial architecture in vivo. J Cell Biol. 2016 Aug 1;214(3):3. doi:10.1083/jcb.201603039 PubMed PMID: 27458135; PubMed Central PMCID: PMC4970326.

90. Montava-Garriga L, Singh F, Ball G, Ganley IG. Semi-automated quantitation of mitophagy in cells and tissues. Mech Ageing Dev. 2020 Jan;185:111196. doi:10.1016/j.mad.2019.111196

91. Jassim AH, Inman DM. Evidence of Hypoxic Glial Cells in a Model of Ocular Hypertension. Invest Ophthalmol Vis Sci. 2019 Jan;60(1):1–15. doi:10.1167/iovs.18-24977 PubMed PMID: 30601926; PubMed Central PMCID: PMC6322635.

92. Dai Y, Hu X, Sun X. Overexpression of parkin protects retinal ganglion cells in experimental glaucoma. Cell Death Dis. 2018 Jan 24;9(2):2. doi:10.1038/s41419-017-0146-9

93. Coughlin L, Morrison RS, Horner PJ, Inman DM. Mitochondrial Morphology Differences and Mitophagy Deficit in Murine Glaucomatous Optic Nerve. Invest Ophthalmol Vis Sci. 2015 Mar;56(3):3. doi:10.1167/iovs.14-16126 PubMed PMID: 25655803; PubMed Central PMCID: PMC4347310.

94. Scorrano L, Ashiya M, Buttle K, Weiler S, Oakes SA, Mannella CA, et al. A distinct pathway remodels mitochondrial cristae and mobilizes cytochrome c during apoptosis. Dev Cell. 2002 Jan;2(1):55–67. doi:10.1016/s1534-5807(01)00116-2 PubMed PMID: 11782314.

95. Hackenbrock CR. Ultrastructural bases for metabolically linked mechanical activity in mitochondria. I. Reversible ultrastructural changes with change in metabolic steady state in isolated liver mitochondria. J Cell Biol. 1966 Aug;30(2):269–97. doi:10.1083/jcb.30.2.269 PubMed PMID: 5968972; PubMed Central PMCID: PMC2107001.

96. Schindelin J, Arganda-Carreras I, Frise E, Kaynig V, Longair M, Pietzsch T, et al. Fiji: an open-source platform for biological-image analysis. Nat Methods. 2012 Jul;9(7):676– 82. doi:10.1038/nmeth.2019

97. Wickham H, Averick M, Bryan J, Chang W, McGowan L, François R, et al. Welcome to the Tidyverse. J Open Source Softw. 2019 Nov 21;4(43):1686. doi:10.21105/joss.01686

98. Barret Schloerke, Di Cook, Joseph Larmarange, Francois Briatte, Moritz Marbach, Edwin Thoen, et al. GGally: Extension of “ggplot2” [R package]. 2025. Available from: https://ggobi.github.io/ggally/

99. Stankowska DL, Minton AZ, Rutledge MA, Mueller BH, Phatak NR, He S, et al. Neuroprotective Effects of Transcription Factor Brn3b in an Ocular Hypertension Rat Model of Glaucoma. Invest Ophthalmol Vis Sci. 2015 Feb;56(2):2. doi:10.1167/iovs.14-15008 PubMed PMID: 25587060; PubMed Central PMCID: PMC4321399.

100. Hu X, Dai Y, Zhang R, Shang K, Sun X. Overexpression of Optic Atrophy Type 1 Protects Retinal Ganglion Cells and Upregulates Parkin Expression in Experimental Glaucoma. Front Mol Neurosci. 2018 Sep 28;11:350. doi:10.3389/fnmol.2018.00350 PubMed PMID: 30323741; PubMed Central PMCID: PMC6172338.

101. Kodati B, Stankowska DL, Krishnamoorthy VR, Krishnamoorthy RR. Involvement of c-Jun N-terminal kinase 2 (JNK2) in Endothelin-1 (ET-1) Mediated Neurodegeneration of Retinal Ganglion Cells. Invest Ophthalmol Vis Sci. 2021 May 12;62(6):13. doi:10.1167/iovs.62.6.13 PubMed PMID: 33978676; PubMed Central PMCID: PMC8131991.

102. Yu J, Luo Y, Lin J, Li Z, Fang Z, He H, et al. Mitochondrial cristae remodeling: Mechanisms, functions, and pathology. Cell Insight. 2025 Dec 1;4(6):100285. doi:10.1016/j.cellin.2025.100285

103. Liang Y, Li Y, Jiao Q, Wei M, Wang Y, Cui A, et al. Axonal mitophagy in retinal ganglion cells. Cell Commun Signal. 2024 Jul 29;22(1):382. doi:10.1186/s12964-024-01761-0

104. Davis C ha O, Marsh-Armstrong N. Discovery and implications of transcellular mitophagy. Autophagy. 2014 Dec 2;10(12):2383–4. doi:10.4161/15548627.2014.981920

105. Davis C ha O, Kim KY, Bushong EA, Mills EA, Boassa D, Shih T, et al. Transcellular degradation of axonal mitochondria. Proc Natl Acad Sci. 2014 Jul;111(26):9633–8. doi:10.1073/pnas.1404651111

106. Klionsky DJ, Abdel-Aziz AK, Abdelfatah S, Abdellatif M, Abdoli A, Abel S, et al. Guidelines for the use and interpretation of assays for monitoring autophagy (4th edition)1. Autophagy. 2021;17(1):1–382. doi:10.1080/15548627.2020.1797280 PubMed PMID: 33634751; PubMed Central PMCID: PMC7996087.

107. Dunn KW, Kamocka MM, McDonald JH. A practical guide to evaluating colocalization in biological microscopy. Am J Physiol-Cell Physiol. 2011 Apr;300(4):4. doi:10.1152/ajpcell.00462.2010

108. Katayama H, Hama H, Nagasawa K, Kurokawa H, Sugiyama M, Ando R, et al. Visualizing and Modulating Mitophagy for Therapeutic Studies of Neurodegeneration. Cell. 2020 May;181(5):1176–1187.e16. doi:10.1016/j.cell.2020.04.025

109. Tribble JR, Kokkali E, Otmani A, Plastino F, Lardner E, Vohra R, et al. When Is a Control Not a Control? Reactive Microglia Occur Throughout the Control Contralateral Pathway of Retinal Ganglion Cell Projections in Experimental Glaucoma. Transl Vis Sci Technol. 2021 Jan 12;10(1):22. doi:10.1167/tvst.10.1.22

110. Lucas-Ruiz F, Galindo-Romero C, Albaladejo-García V, Vidal-Sanz M, Agudo-Barriuso M. Mechanisms implicated in the contralateral effect in the central nervous system after unilateral injury: focus on the visual system. Neural Regen Res. 2021;16(11):2125. doi:10.4103/1673-5374.310670

111. Johnson GA, Kodati B, Nahomi RB, Pham JH, Krishnamoorthy VR, Phillips NR, et al. Mechanisms contributing to inhibition of retinal ganglion cell death by cell permeable peptain-1 under glaucomatous stress. Cell Death Discov. 2024 Jun 28;10(1):305. doi:10.1038/s41420-024-02070-8

112. Pham JH, Zhang W, Le KTT, Kodati B, Amankwa CE, DebNath B, et al. Hybrid molecule SA-10 and its PLGA nanosuspension protect human and rodent retinal ganglion cells against neuronal injury. BMC Neurosci. 2025 Aug 20;26(1):51. doi:10.1186/s12868-025-00971-7

113. Mazzuca MQ, Khalil RA. Vascular endothelin receptor type B: Structure, function and dysregulation in vascular disease. Biochem Pharmacol. 2012 Jul;84(2):147–62. doi:10.1016/j.bcp.2012.03.020

114. Yamboliev IA, Hruby A, Gerthoffer WT. Endothelin-1 Activates MAP Kinases and c-Jun in Pulmonary Artery Smooth Muscle. Pulm Pharmacol Ther. 1998 Apr;11(2–3):205– 8. doi:10.1006/pupt.1998.0139

115. Araki SI, Haneda M, Togawa M, Kikkawa R. Endothelin-1 activates c-Jun NH2-terminal kinase in mesangial cells. Kidney Int. 1997 Mar;51(3):631–9. doi:10.1038/ki.1997.92

116. Rauh A, Windischhofer W, Kovacevic A, DeVaney T, Huber E, Semlitsch M, et al. Endothelin (ET)-1 and ET-3 promote expression of *c-fos* and *c-jun* in human choriocarcinoma via ET_B_ receptor-mediated G_i_ - and G_q_ -pathways and MAP kinase activation. Br J Pharmacol. 2008 May;154(1):13–24. doi:10.1038/bjp.2008.92

117. Wang J, Ma HY, Krishnamoorthy RR, Yorio T, He S. A feed-forward regulation of endothelin receptors by c-Jun in human non-pigmented ciliary epithelial cells and retinal ganglion cells. PLoS ONE. 2017 Sep 22;12(9):e0185390. doi:10.1371/journal.pone.0185390 PubMed PMID: 28938016; PubMed Central PMCID: PMC5609771.

118. Wang S, Long H, Hou L, Feng B, Ma Z, Wu Y, et al. The mitophagy pathway and its implications in human diseases. Signal Transduct Target Ther. 2023 Aug 16;8(1):304. doi:10.1038/s41392-023-01503-7

119. He YL, Li J, Gong SH, Cheng X, Zhao M, Cao Y, et al. BNIP3 phosphorylation by JNK1/2 promotes mitophagy via enhancing its stability under hypoxia. Cell Death Dis. 2022 Nov 17;13(11):966. doi:10.1038/s41419-022-05418-z

120. Picca A, Faitg J, Auwerx J, Ferrucci L, D’Amico D. Mitophagy in human health, ageing and disease. Nat Metab. 2023 Nov 30;5(12):2047–61. doi:10.1038/s42255-023-00930-8

121. Chen G, Kroemer G, Kepp O. Mitophagy: An Emerging Role in Aging and Age-Associated Diseases. Front Cell Dev Biol. 2020 Mar 26;8:200. doi:10.3389/fcell.2020.00200

122. Kim US, Mahroo OA, Mollon JD, Yu-Wai-Man P. Retinal Ganglion Cells—Diversity of Cell Types and Clinical Relevance. Front Neurol. 2021 May 21;12:661938. doi:10.3389/fneur.2021.661938

123. Rheaume BA, Jereen A, Bolisetty M, Sajid MS, Yang Y, Renna K, et al. Single cell transcriptome profiling of retinal ganglion cells identifies cellular subtypes. Nat Commun. 2018 Jul 17;9(1):2759. doi:10.1038/s41467-018-05134-3

